# Transcriptomic changes predict metabolic alterations in LC3 associated phagocytosis in aged mice

**DOI:** 10.1101/2023.03.14.532586

**Authors:** Anuradha Dhingra, John W Tobias, Nancy J Philp, Kathleen Boesze-Battaglia

**Affiliations:** Department of Basic and Translational Sciences, University of Pennsylvania, Philadelphia, PA 19104, USA; Department of Medicine, Penn Genomics and Sequencing Core, Department of Genetics, University of Pennsylvania, Philadelphia, PA, 19104, USA; Department of Pathology, Anatomy, and Cell Biology, Thomas Jefferson University, Philadelphia, Pennsylvania, PA 19107, USA

**Keywords:** cholesterol/trafficking, cholesterol/metabolism, peroxisomes, transcriptomics, monocarboxylate transporters, LC3-associated phagocytosis (LAP), inflammation, retinal pigment epithelium (RPE), fatty acid metabolism

## Abstract

LC3b (*Map1lc3b*) plays an essential role in canonical autophagy and is one of several components of the autophagy machinery that mediates non-canonical autophagic functions. Phagosomes are often associated with lipidated LC3b, to promote phagosome maturation in a process called LC3-associated phagocytosis (LAP). Specialized phagocytes such as mammary epithelial cells, retinal pigment epithelial (RPE) cells, and sertoli cells utilize LAP for optimal degradation of phagocytosed material, including debris. In the visual system, LAP is critical to maintain retinal function, lipid homeostasis and neuroprotection. In a mouse model of retinal lipid steatosis - mice lacking LC3b (*LC3b*^-/-^), we observed increased lipid deposition, metabolic dysregulation and enhanced inflammation. Herein we present a non-biased approach to determine if loss of LAP mediated processes modulate the expression of various genes related to metabolic homeostasis, lipid handling, and inflammation. A comparison of the RPE transcriptome of WT and *LC3b*^-/-^ mice revealed 1533 DEGs, with ~73% upregulated and 27% down-regulated. Enriched gene ontology (GO) terms included inflammatory response (upregulated DEGs), fatty acid metabolism and vascular transport (downregulated DEGs). Gene set enrichment analysis (GSEA) identified 34 pathways; 28 were upregulated (dominated by inflammation/related pathways) and 6 were downregulated (dominated by metabolic pathways). Analysis of additional gene families identified significant differences for genes in the solute carrier family, RPE signature genes, and genes with potential role in age-related macular degeneration. These data indicate that loss of LC3b induces robust changes in the RPE transcriptome contributing to lipid dysregulation and metabolic imbalance, RPE atrophy, inflammation, and disease pathophysiology.

## 1. Introduction

Autophagy, the activity of “self-eating” is vital for cell homeostasis; this catabolic process degrades old, damaged, or abnormal organelles and other substances in the cell cytoplasm [1]. Macroautophagy (hereafter referred to as autophagy, canonical autophagy (CA)) is evolutionary conserved across eukaryotes and involves formation of a double membrane autophagosome, a specialized structure that engulfs cellular components such as protein aggregates, lipids and damaged organelles [2]. Upstream signals such as nutrient deprivation or stress trigger the activation of a multistep autophagy cascade involving several autophagy related proteins (ATGs) culminating in lipidation of microtubule-associated protein light chain 3 (LC3/ATG8) to phosphatidylethanolamine (PE) during autophagosome biogenesis, a step necessary to target autophagosomes for lysosomal degradation [3–6]. Lipid homeostasis requires specialized autophagy processes to maintain mitochondrial and peroxisome health through mitophagy and pexophagy, respectively [5,7].

A sub-set of the autophagy related proteins are involved in a hybrid autophagy-phagocytosis pathway, termed LC3-associated phagocytosis (LAP), considered a non-cannonical autophagy pathway [8–12]. Professional phagocytes such as macrophages, monocytes and, dendritic cells as well as non-professional/specialized phagocytes such as mammary epithelial cells, retinal pigment epithelial (RPE) cells, and sertoli cells utilize LAP for optimal degradation of phagocytosed material including dead cells/cellular debris, and to recycle nutrients such as vitamin A derivatives [6,11,13–15]. While LAP shares molecular machinery with CA, it is a distinct process; unlike CA, LAP is AMPK–mTORC1–ULK1 independent and unresponsive to nutrient status [8,11,16]. In LAP, a sub-set of autophagy machinery proteins conjugate lipidated LC3 (LC3II) to single membrane phagosome to form a LAPosome followed by degradation in the lysosome [8]. LAP and CA are regulated by RUBCN (Run domain Beclin-1 interacting and cysteine-rich containing protein); this protein promotes LAP, and concomitantly suppresses CA [17,18]. Defects in macrophage LAP lead to enhanced pro-inflammatory cytokine production in chronic obstructive pulmonary disease (COPD) and in response to cigarette smoke [10,19]. *in vitro* knockdown of *LC3b* in lung epithelia sensitized these cells to bleomycin-induced apoptosis [20]. Similarly, the *LC3b*^-/-^ mouse, show increased susceptibility to lung injury and fibrosis [20]. LAP mediates anti-inflammatory and immune-suppressive response that in the context of the tumor microenvironment leads to evasion of immune surveillance resulting in tumor progression; and LAP deficiency results in hyper-inflammation [21–23]. Numerous bacteria have evolved strategies to actively evade (or exploit) host cell LAP as a survival mechanism [9,24].

In the eye, LAP supports visual function and lipid homeostasis in the post-mitotic RPE cells. The RPE, residing in close apposition to photoreceptors supplies nutrients to the retina, regulates fluid/ion balance, supports the visual cycle, and maintains the blood retinal barrier [11,25,26]. These functions rely on daily phagocytic clearance of lipid-rich photoreceptor outer segment tips over the lifetime of the retina [27]. RPE-LAP requires, melanoregulin (Mreg, an LC3 interacting protein)-dependent recruitment of LC3II for subsequent phagosome maturation and phagolysosome formation leading to degradation of the phagocytosed cargo [12,28]. Some of the degradation by-products are removed from the cell by transport to the choroid and others are recycled; they replenish essential components and metabolites needed by the neural retina [6,11,29]. Mouse models of defective RPE-LAP (*Atg5ΔRPE*, *Mreg*^-/-^ and *LC3b*^-/-^) exhibit increased phagosome accumulation, inadequate recovery of retinoids and reduced visual capacity [11,12,28,30].

RPE is intimately associated with the neural retina and RPE-LAP plays an important role in lipid regulation in the eye. Indeed, several studies have demonstrated that lipid dysregulation contributes to the pathogenesis of age-related macular degeneration (AMD) [31–34]. The RPE exploits LAP to efficiently degrade lipid and protein rich OS material daily; it metabolizes OS lipids to generate β-hydroxybutyrate (by ketogenesis), which may support energy needs in the outer retina [35,36]. Abrogation of LAP, defective processing of lipid rich phagosomes in *LC3b*^-/-^ RPE causes phagosome accumulation resulting in decrease in fatty acid oxidation and ketogenesis as well as disrupted lipid homeostasis. This results in an increase in RPE and sub-RPE lipid deposits, elevated levels of lipid peroxidation products, pro-inflammatory sterol, 7-ketocholesterol (7KCh) and decreased levels of lipid pro-survival factors Neuroprotectin D1 (NPD1) and Maresin-1 [28]. Moreover, these mice show sub-retinal recruitment of microglia [28]. These studies point to a vital role for LAP in lipid regulation with a loss of lipid/cholesterol homeostasis contributing to an AMD-like phenotype [11,12,28,30]. Over time in aging *LC3b*^-/-^ mice, a chronic low–grade inflammatory response (para-inflammation) is manifested which combined with observed lipid steatosis is reminiscent of non-alcoholic fatty liver disease [37].

Herein, we investigate how loss of LC3b affects gene expression using an unbiased, RNAseq based comparative transcriptomics approach, thus allowing us to further uncover the mechanistic aspects of defective LAP leading to inflammation and lipid metabolic dysregulation. We provide a detailed report of the differentially expressed genes (DEGs) and perform gene ontology-based enrichment analyses and hallmark pathway analyses to decipher affected pathways. Our results show that in the *LC3b*^-/-^ mice, inflammatory response, cholesterol homeostasis, and complement activation are upregulated, while fatty acid metabolism and oxidative phosphorylation are downregulated. Moreover, the data provides novel insights into gene expression changes in the solute carrier family and some RPE signature genes suggesting metabolic changes resulting from defective LAP. Finally, we identify 8 DEGs that have corresponding risk alleles reported for AMD in a large genome-wide association study (GWAS) [38] and experimentally validate the correlation between gene transcription (based on RNAseq) and protein expression for one of these DEGs. These results point to molecular links between defective LAP and AMD-like pathology and potentially can be extended to other age-related pathologies.

## 2. Results

### 2.1. RNAseq data source and quality (High Throughput RNA Sequencing Data Set Preparation for Analysis)

Aged *LC3b*^-/-^ mice show retinal lipid imbalance and enhanced pro-inflammatory micro-environment. This is due to RPE and sub-RPE lipid deposits and a concomitant decrease in the synthesis of protective lipid species, NPD1 and maresin-1 [28]. To identify RPE specific changes in the *LC3b*^-/-^ mouse at the gene expression level, we prepared RNA from RPE isolated from mouse retina. The experimental design used for RPE cell isolation for subsequent RNA preparation and sequencing is illustrated in Figure 1A. Sequencing was performed using Illumina NovaSeq 6000 SP flow cell with the XP workflow (100 bp single read sequencing) to a minimum depth of 28.54MR with a median depth of 30.4 MR (+/- 5.8 MR). Total reads for the samples ranged from 26,704,479 to 35,616,153. For each sample, over 85% of reads mapped to the transcriptome. Principal component analysis (PCA) shows the overall variation among the samples with a clear separation in the first two components

**Figure 1.**
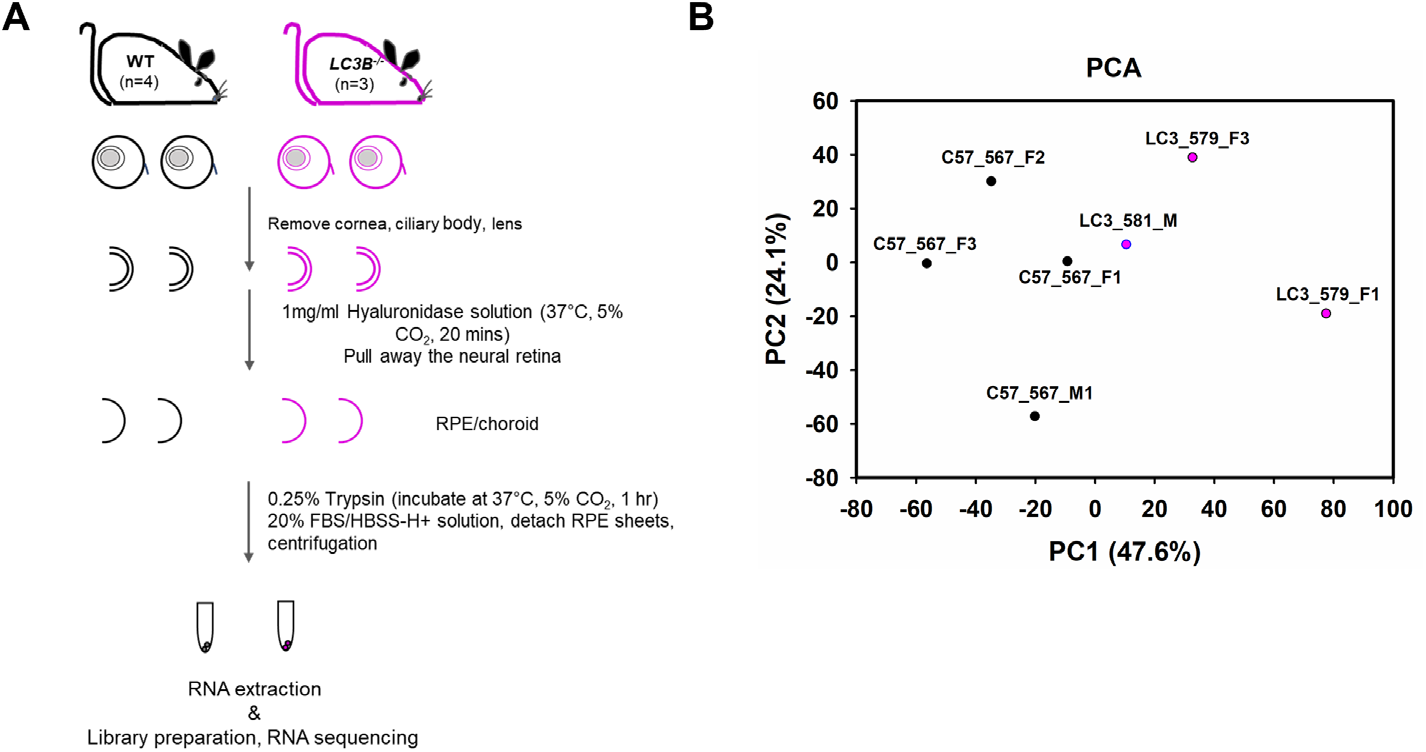
RPE cell isolation for RNAseq and Principal component analysis. (A) Experimental design to isolate RPE from mouse eyes. ~24-month old WT and *LC3b*^-/-^ mice were sacrificed, eyeballs were removed and processed to isolate RPE cells by enzymatic treatments as indicated. Total RNA was extracted and used for library preparation and RNA sequencing. (B) Principal component analysis (PCA), X and Y axis show PC1 and PC2, respectively.

### 2.2. RNA-Sequencing analysis to identify differentially expressed genes (DEGs)

Differential expression between *LC3b*^-/-^ and WT was analyzed using DESeq2. Genes showing fold change of ≤ −1.5 or ≥ 1.5, and an adjusted p-value < 0.05 were considered differentially expressed. There were 1533 DEGs between *LC3b*^-/-^ and WT, 1118 (~73%) up-regulated and 415 (27%) downregulated (Figure 2, Volcano plot). A list of DEGs identified in this dataset along with their Log2 fold change and adjusted p-value is provided in Table S1. As expected, Map1lc3b was one of the most highly downregulated genes by adjusted p-value; furthermore, there was no compensatory upregulation of a related isoform *Map1lc3a* (gene encoding for LC3a) or other members of the ATG8/LC3 family (*Gabarap*, *Gabarapl1*, *Gabarapl2*) in the *LC3b*^-/-^ mouse (Figure 2) [5,28,39–41].

**Figure 2.**
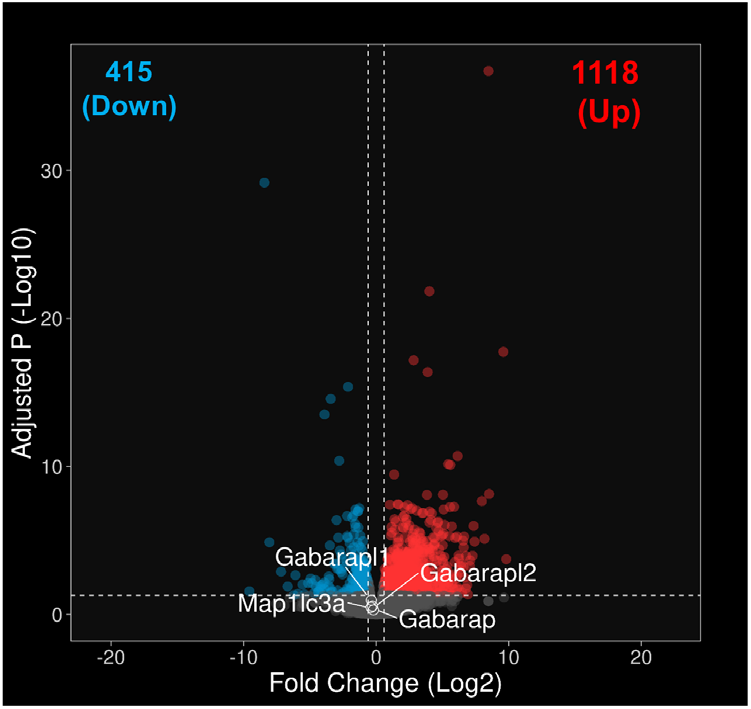
Differentially Expressed Genes between the *LC3b*^-/-^ and WT RPE. Volcano plot of DEGs between the *LC3b*^-/-^ and WT RPE with −Log10 of the adjusted P value on the Y axis and Log2 of the fold change expression on the X axis. Each point represents a single gene. DEGs were identified by using a cut-off of Padj < 0.05 and fold change of ≤ −1.5 or ≥ 1.5 and have been shown in red (upregulated DEGs), cyan (dowregulated DEGs), or grey dots (unchanged genes). Genes that are upregulated in the *LC3b*^-/-^ are on the right (Up) and downregulated genes are on the left (Down). Genes belonging to ATG8/LC3 family (except for *Map1lc3b*) are indicated. *Map1lc3b* with adjusted P (-Log10) of 168.4 is not shown in the volcano plot.

### 2.3. RNA-Sequencing reveals pathways related to inflammation and metabolism

We performed Gene Ontology (GO) based enrichment analysis using the Metascape tool under 3 categories: Molecular Functions (MF), Cellular components (CC), and Biological Processes (BP). Table 1 shows a list of GO terms for top 10 DEGs under each of these categories. Closer examination of these GO terms revealed that the majority overlap with upregulated DEGs. This is not surprising since there are many more upregulated DEGs (>2.5 fold) in our list. Next, we performed a similar enrichment analysis across all upregulated and downregulated genes separately. Figure 3A shows top 5 most enriched GO terms for each of the 3 categories; more than 5 GO terms have been shown in cases where LogP values were exactly same for more than one term. Interestingly, some of the most enriched GO terms for the upregulated genes are related to inflammation (Inflammatory response and cellular response to cytokine stimulus) and cell-cell adhesion (Figure 3A), whereas the downregulated DEGs were enriched in GO terms related to metabolism including fatty acid metabolic process, transmembrane transporters, and mitochondria (Figure 3B). There was no significant enrichment of GO terms corresponding to phagosome maturation (GO:0090382) or phagosome lysosome fusion (GO:0090385). Reactome pathway analysis also supported these results; interleukin and cytokine signaling seen in 2 of the top 5 upregulated DEGs (Figure 3C) while, metabolism of lipids and fatty acid metabolism in the downregulated DEGs (Figure 3D).

**Table 1.**
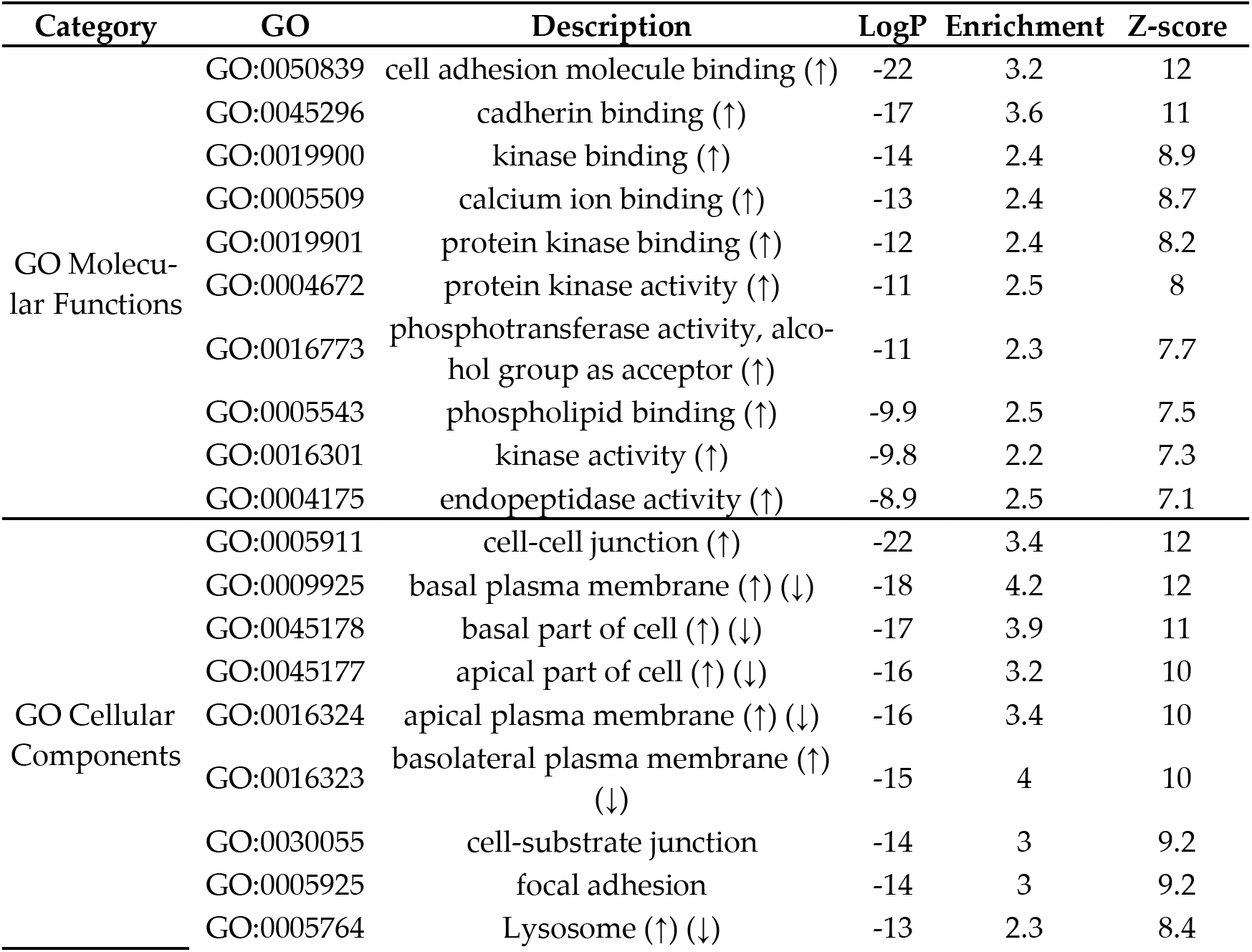
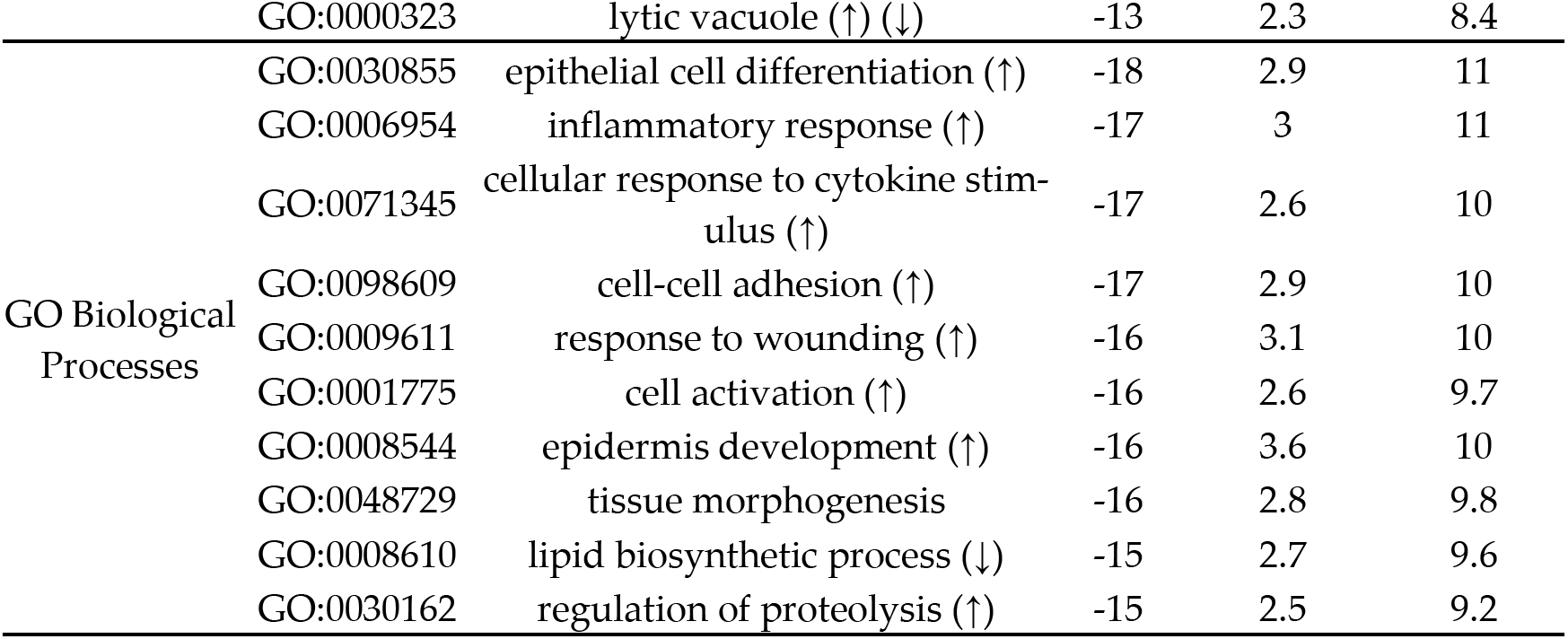
Top10 most enriched Gene ontology terms (GO) (selected based on the p-values) across all DEGs. GO terms with up (↑) or down (↓) arrows indicate those enriched with upregulated or downregulated genes, respectively. Analysis was performed using Metascape.

**Figure 3.**
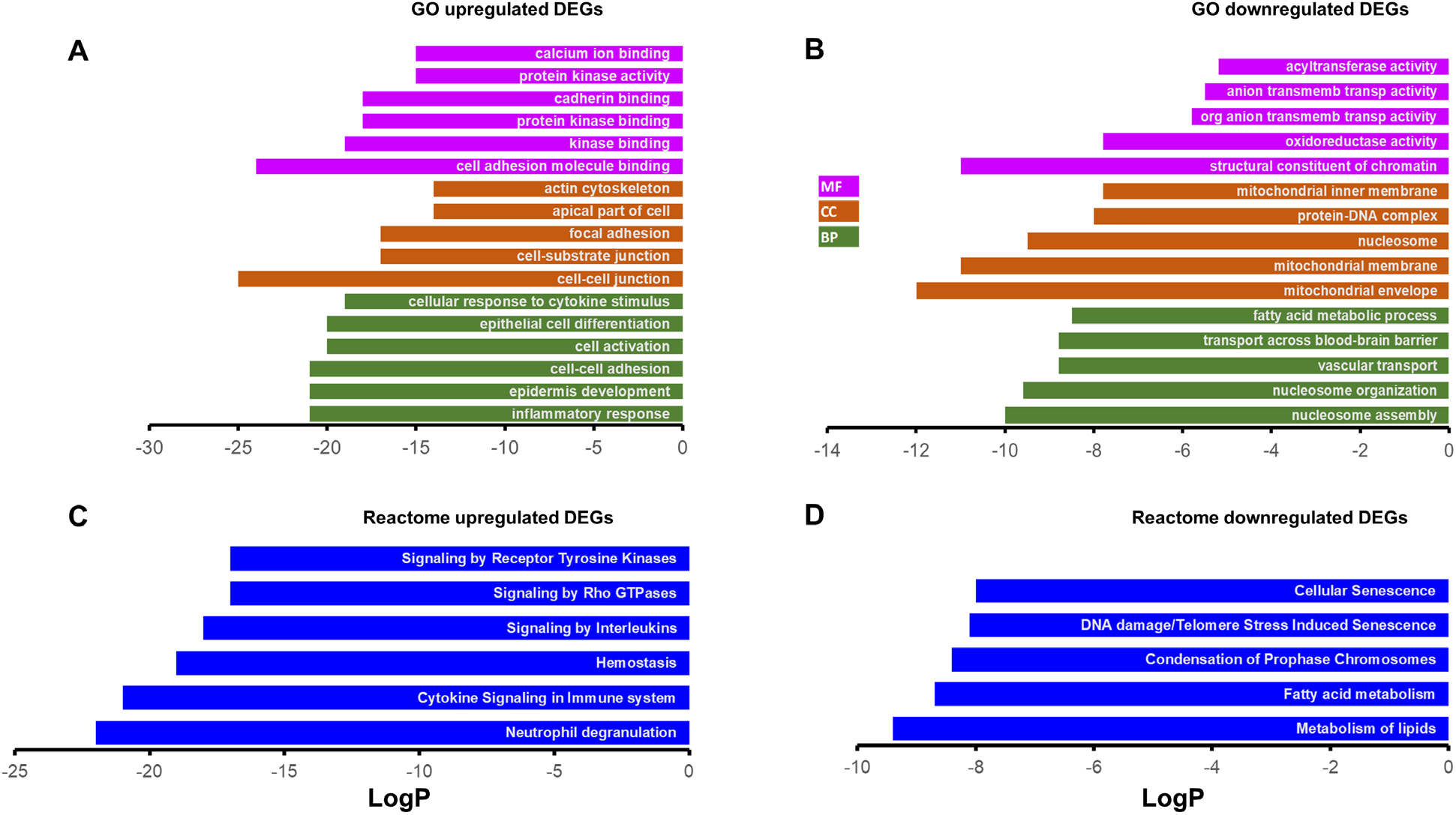
Gene Ontology (GO) and reactome based gene enrichments. The top 5 most enriched GO terms of upregulated DEGs (A) and downregulated DEGs (B). The top 5 most enriched reactome gene terms for upregulated DEGs (C) and downregulated DEGs (D). The terms were selected based on the lowest LogP values (>5 GO terms shown in cases where LogP values were exactly same for more than one term). Analysis was performed using Metascape. MF: molecular function; CC: cellular component; BP: biological process.

Also, an unfiltered list of genes ranked by their DESeq2 statistic was analyzed with GSEA against the hallmark gene collections in the Molecular Signatures Database (MSigDB). GSEA identified 34 significantly different (FDR q-val < 0.05) hallmark pathways; 28 were upregulated (Figure 4, top) and 6 were downregulated (Figure 4, bottom). Upregulated pathways were dominated by inflammation/related pathways (eg, inflammatory response, complement, cholesterol homeostasis); and those enriched by down-regulated genes were dominated by metabolic pathways (eg., oxidative phosphorylation, fatty acid metabolism, and peroxisome). We have previously established that LC3b plays a critical role in lipid mediated homeostasis in the RPE; loss of LC3b leads to decrease in fatty acid oxidation, peroxisomal turnover defects, age-dependent lipid accumulation, oxidative stress, and pro-inflammatory microenvironment [28,42].

**Figure 4.**
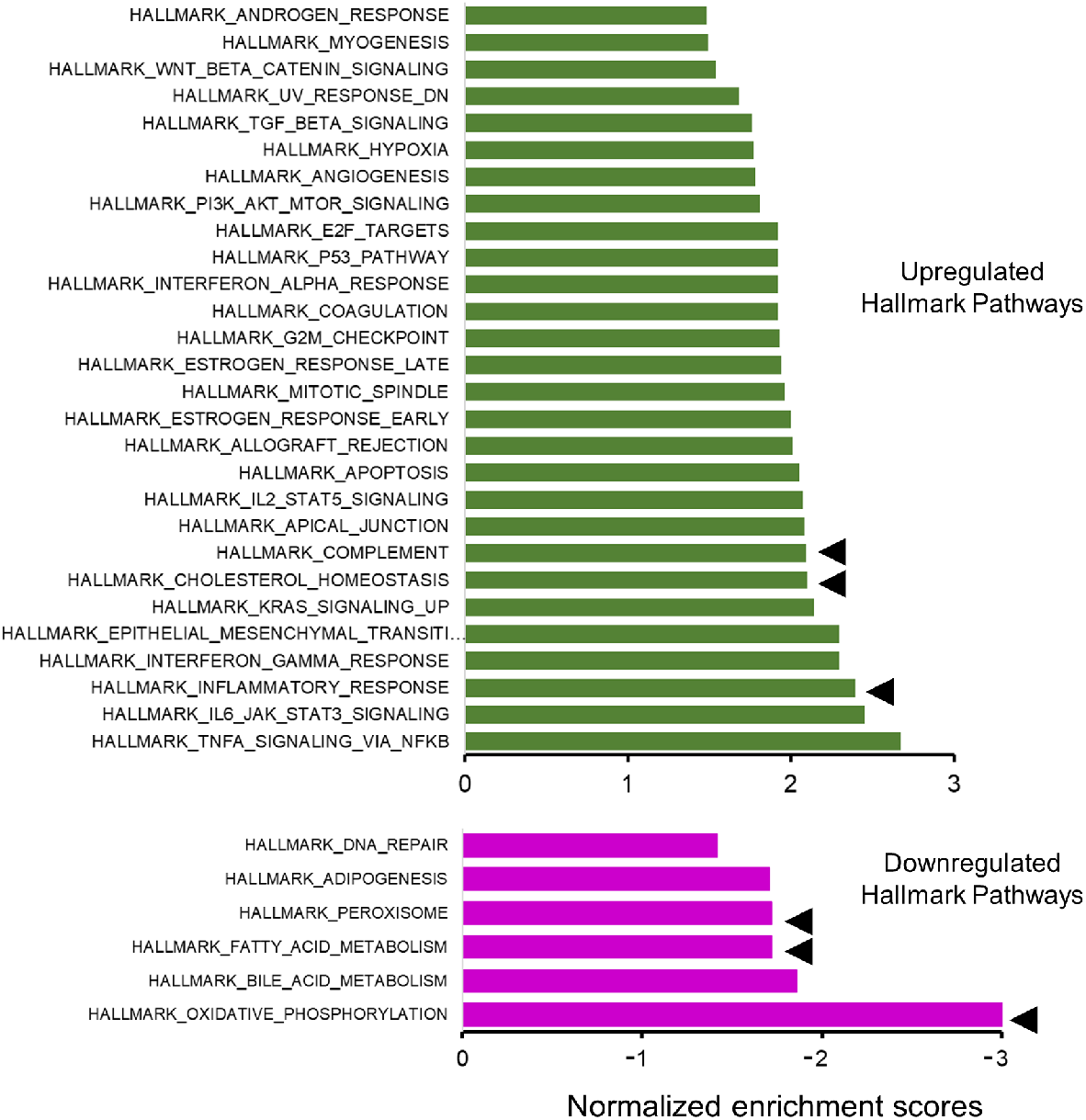
Gene set Enrichment Analysis (GSEA). Bar chart showing absolute normalized enrichment scores for the hallmark pathways that were significantly different between the *LC3b-/-* and WT (false discovery rate or FDR <0.05); GSEA identified 34 significantly different hallmark pathways, 28 were upregulated (top panel) and 6 were downregulated (bottom panel). Pathways indicated by arrowhead (◄) were selected for further analysis.

*LC3b*^-/-^-RPE exhibits decrease in fatty acid oxidation and ketogenesis along with chronic low-grade inflammation [28]. To understand the contribution of gene expression changes associated with lipid metabolic imbalance and pro-inflammatory environment, we further examined some of the key hallmark pathways in relation to RPE physiology and metabolism (Figure 4, pathways marked by arrowheads). The hallmark inflammatory response gene set is comprised of 196 genes; 112 genes were part of the leading edge with a normalized enrichment score (NES) of 2.39 (Figure 5A, S1A). The hallmark complement gene set includes 187 genes; 85 genes were part of the leading edge with an NES of 2.1 (Figure 5B, S1B). The hallmark cholesterol homeostasis gene set includes 74 genes; 38 genes were part of the leading edge with an NES of 2.1 (Figure 5C, S1C).

**Figure 5.**
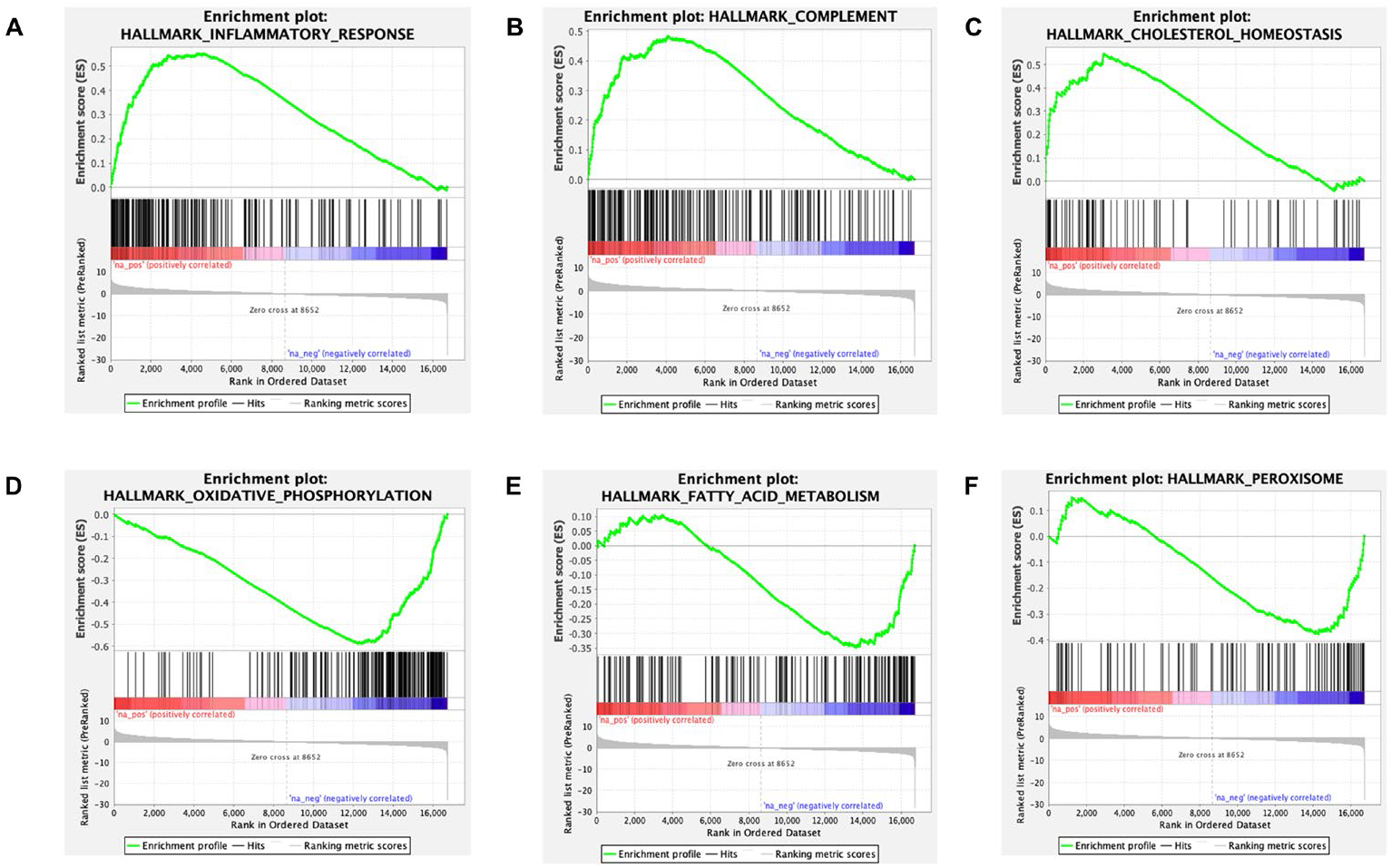
GSEA enrichment profile of upregulated and downregulated pathways. (A) A GSEA enrichment plot of the hallmark inflammatory response pathway with a normalized enrichment score 2.39 and an enrichment score of 0.55 indicating a positive relation. The leading edge includes 112 genes. (B) A GSEA enrichment plot of the hallmark complement pathway with a normalized enrichment score 2.1 and an enrichment score of 0.48 indicating a positive relation. The leading edge comprised of 85 genes. (C) A GSEA enrichment plot of the hallmark cholesterol homeostasis pathway with a normalized enrichment score 2.1 and an enrichment score of 0.55 indicating a positive relation. The leading edge comprised of 38 genes. (D) A GSEA enrichment plot of the hallmark oxidative phosphorylation pathway with a normalized enrichment score −3.02 and an enrichment score of −0.59 indicating a negative relation. The leading edge comprised of 138 genes. (E) A GSEA enrichment plot of the hallmark fatty acid metabolism pathway with a normalized enrichment score −1.72 and an enrichment score of −0.35 indicating a negative relation. The leading edge comprised of 57 genes. (F) A GSEA enrichment plot of the hallmark peroxisome pathway with a normalized enrichment score −1.72 and an enrichment score of −0.38 indicating a negative relationship. The leading edge comprised of 35 genes.

The hallmark oxidative phosphorylation gene set is comprised of 199 genes; 138 genes were part of the leading edge with an NES of −3.02 (Figure 5D, S2A). The hallmark fatty acid metabolism gene set includes 155 genes; 57 genes were part of the leading edge with an NES of −1.72 (Figure 5E, S2B). The hallmark peroxisome gene set is comprised of 102 genes; 35 genes were part of the leading edge with an NES of −1.72 (Figure 5F, S2C).

### 2.4. Several solute carrier family genes are differentially expressed

Transporters play one of the most essential roles in metabolic homeostasis [43]. Anion transmembrane transporter activity was identified as one of the top GO terms in the pathway enrichment analysis of downregulated DEGs (Figure 3B). As RPE is intimately connected with choroid (basal), and neural retina (apical) and has an important regulatory role in movement of metabolites and ions into and out of retina, we broadened our analysis to look specifically at solute carrier family gene expression. There were 43 DEGs (depicted in the heat map, Figure 6A top) between *LC3b*^-/-^ and WT out of a total of 390 solute carrier genes (χ^2^=9.599; two-tailed P value <0.05). Interestingly, *Slc16a1* and *Slc16a8* encoding for MCT1 and MCT3 respectively, two of the key lactate transporter genes expressed in the RPE [44,45] are downregulated; while *Slc16a11* (encoding for MCT11, an orphan transporter) expression is upregulated in *LC3b*^-/-^ RPE [46]. Moreover, expression of *Bsg,* encoding for Basigin (CD147), a protein required for proper trafficking of MCT1 and MCT3 to the plasma membrane, was also downregulated (Table S1, Figure 6A, bottom) [47,48]. Consistent with the transcriptomic analysis, we observed ~40% decrease in MCT3 protein levels in the *LC3b*^-/-^ compared to the WT control (Figure 6B). Mouse RPE/choroid flat mounts also showed a decrease in MCT3 with a mis-localization of much of the remaining MCT3 as intracellular and not plasma membrane associated, consistent with the decrease in *Bsg* (Figure 6C).

**Figure 6.**
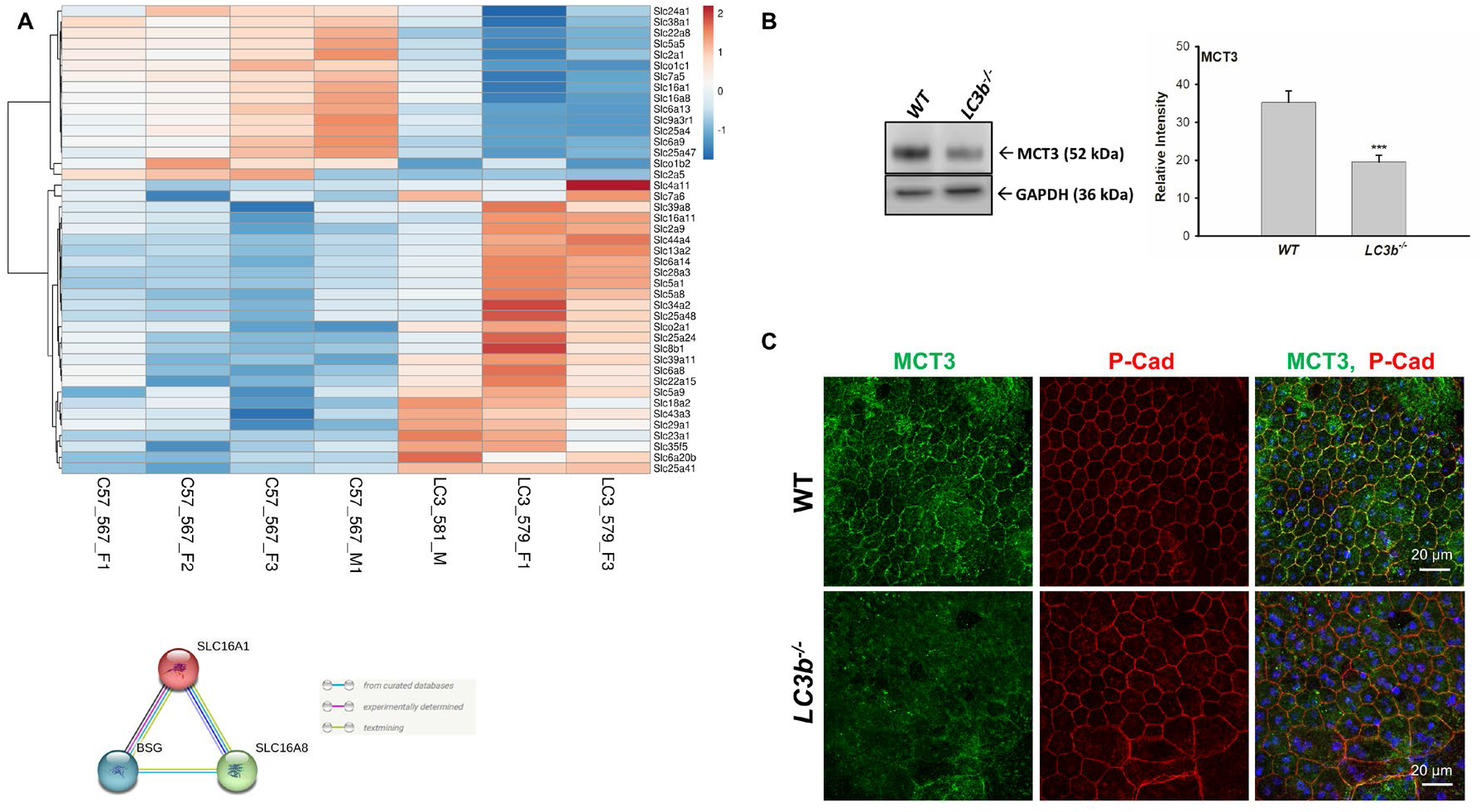
Analysis of the solute carrier family genes. (A) Top: Heat map of various solute carrier family DEGs. The variance stabilized counts were converted to heat map colors using Clustvis software. Rows are centered; unit variance scaling is applied to rows. Rows are clustered using correlation distance and average linkage. The intensity scale ranges from low expression (dark blue) to high expression (red) relative to the median of the gene’s expression across all samples. Gene names (left) and mouse ID (bottom) are indicated. χ2 = 9.599 with 1 degrees of freedom, two-tailed p-value=0.0019. Bottom: String diagram showing nexus between Slc16a1, Slc16a8, and Basigin (Bsg). (B) Left: Western blots of RPE/choroid from WT and *LC3b*^-/-^. Immunoblot analysis was performed with antibodies against MCT3 and loading control (GAPDH). Right: Quantification of mean intensity ±SEM of MCT3 relative to GAPDH (N=3 eyes). (C) Representative confocal images of RPE/choroid flat mount from WT (top) and *LC3b*^-/-^ (bottom) immuno-stained for MCT3 (green) and P-cadherin (red). Images are projections from 5 μm stacks captured using 60X water objective (NA1.2) on Nikon A1R laser scanning confocal microscope.

### 2.5. Loss of LC3b is associated with differential expression of select RPE signature and lipid metabolism related genes

Gene expression alterations in some of the key lactate transporter genes (such as *Slc16a1*) prompted us to evaluate expression of RPE signature and lipid metabolism related genes. We used mouse orthologs from a list of RPE signature genes (human) reported previously [49]. Our analysis revealed significantly different expression profile between *LC3b*^-/-^ and WT (χ^2^= 6.43; two-tailed P value <0.05) with 19 DEGs among 149 genes in the list. Expression profile of the DEGs is depicted in the heat map (Figure 7A). Gene expression of *Rbp1* (retinoid binding protein1), *Enpp2* (Ectonucleotide Pyrophosphatase/Phosphodiesterase 2), and *Ptgds* (Prostaglandin D2 Synthase), further suggest some functional decline in the *LC3b*^-/-^ RPE. Moreover, expression of *Slc6a20b* (Sit1, a proline and betaine transporter) is upregulated (Figure 7A). Interestingly, expression of *Best1* and *Rpe65* two well studied RPE markers did not change significantly (Figure S3A and B) [50].

**Figure 7.**
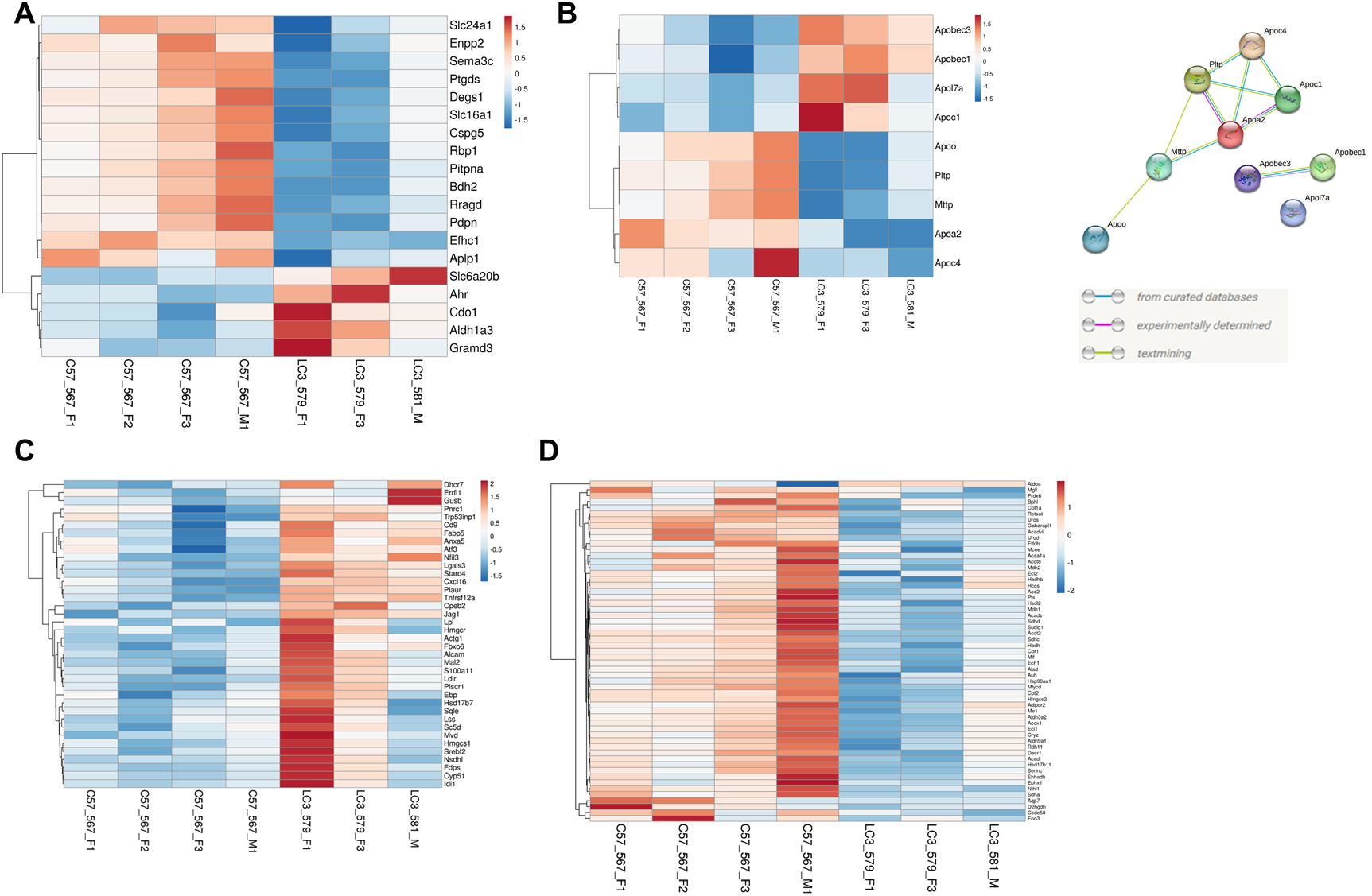
Heat maps depicting DEGs or leading-edge genes in different gene sets. (A) Heat map of the DEGs between the *LC3b*^-/-^ and WT from RPE signature gene set. 19 DEGs out of 149 RPE signature genes examined; χ^2^ = 6.43 with 1 degrees of freedom (two-tailed p-value=0.0112). (B) Heat map of the DEGs (between the *LC3b*^-/-^ and WT RPE) overlapping with lipoprotein metabolism associated genes. 9 DEGs out of 41 lipoprotein metabolism associated genes examined; χ^2^ = 13.93 with 1 degrees of freedom (two-tailed p-value = 0.0002) (C) Genes in the leading edge of cholesterol homeostasis hallmark pathway (upregulated). (D) Genes in the leading edge of fatty acid metabolism hallmark pathway (downregulated). For all the heatmaps, the variance stabilized counts were converted to colors using Clustvis software. Rows are centered; unit variance scaling is applied to rows. Rows are clustered using correlation distance and average linkage. The intensity scale ranges from low expression (dark blue) to high expression (red) relative to the median of the gene’s expression across all samples. Gene names (left) and mouse ID (bottom) are indicated.

On a molecular scale our GSEA analysis showed transcriptional changes in fatty acid metabolism and cholesterol homeostasis. We sought to further explore if LAP mediated processes modulate the level of genes related to lipid transport/handling. Gene expression profiles for lipoprotein metabolism associated genes including apolipoproteins (30 genes), apolipoprotein B mRNA editing enzyme catalytic polypeptides 1-4 (4 genes), *Apobr, Mttp, Pltp*, and *Lcat*, *Acat1*, *Acat2*, and *Lrp8* also showed significance differences between *LC3b*^-/-^ and WT with 9 DEGs out of 41 genes examined (χ^2^=13.93; two-tailed P value <0.05) (Figure 7B, left). PPI string diagram (Figure 7B, right) depicting interaction between the differentially expressed genes - *Apoc4*, *Apoc1*, *Apoa2*, *Pltp*, and *Mttp* suggests altered lipoprotein metabolism in the *LC3b*^-/-^ RPE with *Apoa2* potentially playing a central role.

Lipoprotein metabolism is intimately linked to cholesterol biosynthesis and transport. GSEA of cholesterol homeostasis pathways show upregulation of genes for cholesterol biosynthesis (*Hmgcr*, *Hmgcs1*, *Fdps*, *Sqle*, *Cyp51a1*, *Idl1*, *Nsdhl* and others), as well as genes associated with cholesterol trafficking/import (*Stard4*, *Ldlr*) [51]. Core enrichment subset in the cholesterol homeostasis pathway also includes genes coding for proteins involved in lipid metabolic disorders some of which link lipoprotein metabolism with cholesterol homeostasis (*Lpl*, *Ldlr*) and the cholesterol biosynthetic enzymes, *Dhcr7*, *Nsdhl*, and *Ebp*) (depicted as heat map in Figure 7C) [52,53]. GSEA of fatty acid metabolism pathway shows downregulation of genes involved in ketogenesis (*Hmgcs2*), several genes critical for the TCA cycle (*Sdhc*, *Sdha*, *Sdhd*, *Aco2*, *Suclg1*), peroxisomal fatty acid oxidation (Acox1, Acaa1), and mitochondrial β-oxidation (*Acadl*, *Acadvl*, *Cpt2*, *Cpt1a*) (depicted as heat map in Figure 7D). Acox1 catalyzes desaturation of acyl-CoA esters which is the initial committed step in peroxisomal fatty acid oxidation [54,55].

### 2.6. DEGs with potential role in age-related retinal diseases

Dysregulation in lipid/cholesterol homeostasis is linked to age-related diseases and loss of RPE function [56–58]. Several features observed in the *LC3b*^-/-^ RPE such as reduced phagocytosis, lipid deposits, recruitment of immune cells, and complement pathway activation are reminiscent of AMD-like pathophysiology [28,59–61]. To gain insight into the molecular changes underlying this pathology, we analyzed expression of genes with potential role in AMD.

We started with a list of 41 genes based on a list identified as genes with top priority and statistical significance combined in a genome-wide association study (GWAS) published by the International AMD Genomics Consortium (IAMDGC). Currently, this is both the most recent and largest AMD GWAS study and reported 52 independent genetic signals distributed over 34 loci associated with AMD at genome-wide significance [38,62]. There was significantly different expression profile between *LC3b*^-/-^ and WT (χ^2^= 12.02; two-tailed P value = 0.0005) with 8 DEGs out of 37 mouse orthologs identified (depicted in the heat map, Figure 8) including *Rdh5*, *Slc16a8*, and *Htra1*. Slc16a8 (MCT3) plays an important role in regulating ionic composition of outer retina and *Slc16a8*^-/-^ mice showed reduced visual function [63]. A subset of patients with *RDH5* (retinol dehydrogenase 5) mutation develop macular atrophy; and reduced RDH5 activity is considered as a risk factor for AMD [64,65]. Increased protein expression of *HTRA1* (High Temperature Requirement A Serine Peptidase 1), a serine protease was seen in iPSC-derived RPE cell line from subjects carrying AMD risk-associated 10q26 locus [66].

**Figure 8.**
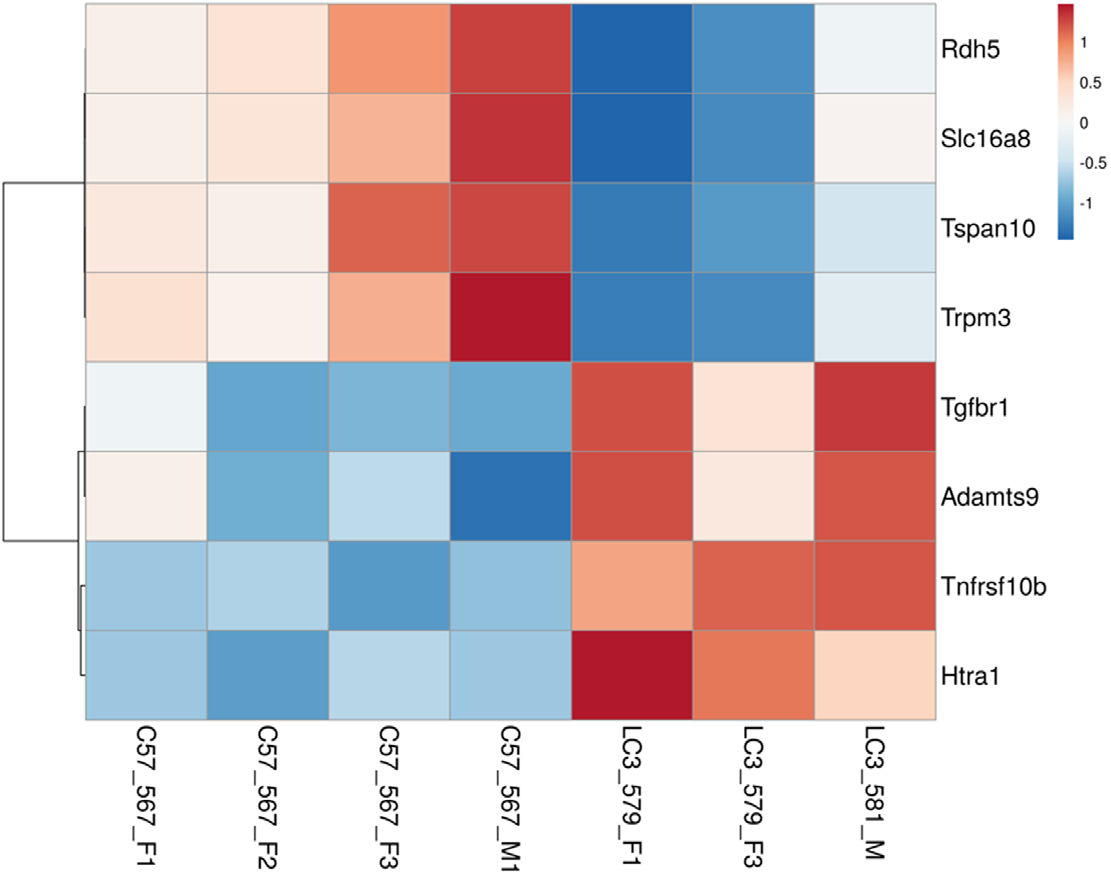
Heat map of the DEGs overlapping with AMD loci genes. AMD loci genes with top priority based on biological and statistical evidence combined (from Fritsche et. al. 2016) were used for the analysis. The variance stabilized counts were converted to heat map colors using Clustvis software. Rows are centered; unit variance scaling is applied to rows. Rows are clustered using correlation distance and average linkage. The intensity scale ranges from low expression (dark blue) to high expression (red) relative to the median of the gene’s expression across all samples. Gene names (left) and mouse ID (bottom) are indicated. χ^2^ equals 12.02 with 1 degrees of freedom; two-tailed P value = 0.0005.

## 3. Discussion

Post-mitotic cells such as neurons, RPE cells, and cardiac myocytes utilize autophagic processes to sustain homeostasis; these cells get rid of unwanted material (waste) and recycle cellular components thus maintaining intracellular quality control over their lifetime [6,67]. In the RPE, LAP plays a necessary role in cell homeostasis through optimal clearance of phagocytosed OS fragments; this supports 1) visual cycle by recycling retinoids, 2) retinal metabolism, eg by metabolizing lipids to generate ketones and thus providing energy to the neural retina, and by producing insulin locally to support homeostasis, 3) RPE and retinal health and function by preventing buildup of excess lipids, 4) ensuring the synthesis of anti-inflammatory lipids [11,28,30,43,68]. LAP thus serves a protective role in the retina. We have previously established that the LC3b, an LC3 isoform, plays a crucial role in RPE-LAP; it is critical for lipid homeostasis and in regulating the inflammatory state of the retina/RPE [5,28,41]. Here, using RNAseq based comparative transcriptomics, we identified 1533 DEGs with ~73% of these upregulated and 27% downregulated in the *LC3b*^-/-^ RPE. There was no compensatory upregulation of any other Atg8/LC3 family members - *LC3a*, *Gabarap*, *Gabarapl1*, and *Gabarapl2*, consistent with previous studies showing no upregulation of *LC3a* in *LC3b*^-/-^ [5,28,39,41,69]. In the absence of LC3b, inflammatory and complement related pathways are upregulated while fatty acid metabolism, oxidative phosphorylation, and peroxisome pathways are downregulated. To the best of our knowledge, this is the first report using RNAseq based expression profiling in a defective LAP model.

### 3.1. LC3b and lipid metabolic dysregulation

Daily phagocytosis of OS presents a unique set of challenges for the RPE; estimates suggest that each RPE processes ~0.08-0.15 pmoles of OS derived fatty acid per day [6,27,35]. Excessive lipid accumulation and formation of lipid-peroxidation adducts in RPE/Ch is a common feature of mouse models of defective phagosome transport and degradation [30,57,70]. The RPE utilizes LAP in the daily ingestion of lipids that it degrades to provide metabolic intermediates; the lipids provide a critical energy source for the RPE through fatty acid oxidation and metabolically couple with photoreceptors to provide β-hydroxybutyrate (β-HB) [35,36,42].

Herein, we found that the hallmark pathways for oxidative phosphorylation and fatty acid metabolism are significantly downregulated in 24-month-old old *LC3b*^-/-^ mice. Leading-edge gene subsets of these pathways include *Acadvl*, *Cpt2*, *Cpt1a*, and *Acadl* (involved in mitochondrial fatty acid β-oxidation), *Sdhc*, *Sdha*, *Sdhd*, *Aco2*, *Suclg1* (involved in TCA cycle), and *Hmgcs2* (rate limiting enzyme in ketogenesis). Approximately 30% of OS lipids are very long chain fatty acids (VLCFAs, C ≥ 20); which are preferentially catabolized in peroxisomes via β-oxidation [55,71]. Therefore, efficient lipid catabolism in the RPE requires coordination of β-oxidation pathways within both mitochondria and peroxisomes [42,55,71,72]. Our GSEA analysis showed significant downregulation of peroxisome pathways, both biogenesis and turnover, as indicated by alterations in *Acox1*, and *Cat* as well as *Pex6*, *Pex11a* and *Pex2*. The protein products of the Acox gene family, the acyl-CoA oxidases (ACOXs) catalyze desaturation of the AcylCoA esters in the committed step of peroxisome fatty acid oxidation [55,71]. Hydrogen peroxide generated in the ACOX reaction is rapidly degraded by the anti-oxidant enzyme catalase in a coupled reaction. Peroxisomes are the main site for catalase dependent antioxidant activity to maintain redox homeostasis of the RPE [73]. In the RPE, peroxisome turnover is an LC3b dependent process; *LC3b*^-/-^ RPE have elevated peroxisome numbers with reduced antioxidant function reminiscent of aging/disease RPE [42,73,74]. Furthermore, peroxisome catalase activity is lower in primary RPE from aged donors and those with AMD [75]. While the interplay between LAP, peroxisome FAO and peroxisome turnover is a subject of future investigation, downregulation of the hallmark peroxisome pathway in the *LC3b*^-/-^ further lends support to decline in peroxisome function contributing to lipid dysregulation, higher oxidative stress, and disease pathogenesis.

Mitochondrial ketogenesis often serves as a fat disposal conduit, specifically when fatty acid oxidation exceeds mitochondrial capacity [37,76]. Such decreased capability may be due to mitochondrial dysfunction, aging, or disease [77] as well as lipid overload for example after a high fat meal. By disposing of acetyl-CoA, a product of FAO, ketogenesis prevents the need for terminal oxidation through the TCA cycle thus decreasing oxidative stress due to enhanced TCA flux [78]. Progressive mitochondrial dysfunction is a major contributor to the pathophysiology of age related diseases of the outer retina [56,79,80]. Mice lacking *Hmgcs2* exhibited ketogenic deficiency and develop non-alcoholic fatty liver disease spontaneously [81]. *RetSat*, coding for retinol saturase, an oxidoreductase with diverse functions including retinoid metabolism was also included in the leading edge gene subsets. In adipocytes, *RetSat* expression is regulated by PPAR γ and is proposed to be involved in insulin sensitivity [82,83]. *Retsat*^-/-^ mice show reduced production of all-trans-13,14-dihy-droretinol from dietary vitamin A, increased accumulation of neutral lipids and increased adiposity [82,84].

Our transcriptomic analysis provided unique insight into the regulation of metabolic pathways at the level of gene expression in RPE-LAP deficient cells. Although fatty acid metabolism and oxidative phosphorylation pathways were down regulated cholesterol homoeostasis pathways were upregulated. The RPE not only synthesizes cholesterol through the mevalonate pathway but also takes up cholesterol from the circulation as well as the daily phagocytosis of OS tips with cholesterol homeostasis regulated via cholesterol efflux [85–87]. In the aging macula, dysregulation of cholesterol metabolism and lipoprotein processing are considered contributors to the onset and development of age-related degeneration [57]. Non-alcoholic fatty liver disease is similarly associated with dysregulated cholesterol metabolism, and manifests on a molecular level as cholesterol accumulation in the mitochondria [88]. Schirris and colleagues [89] using metabolic network modeling suggest cholesterol biosynthesis as a potential compensatory pathway in mitochondrial dysfunction serving as a means by which to restore NADH and NADPH balance. Our own transcriptomic studies (Figure 5D, S2A) and previous in vitro studies suggest mitochondrial dysfunction, in LAP deficient RPE [90]. If compensatory, cholesterol biosynthesis as a means to maintain homeostasis present a unique challenge to the RPE as cholesterol is also taken up through daily LAP dependent degradation of OS tips, loss of which results in high levels of the pro-inflammatory sterol, 7-keto cholesterol, a component of AMD-associated drusen [91].

A critical aspect of cholesterol efflux in the RPE is cholesterol esterification and subsequent packaging and secretion in lipoprotein-like particles (LLP) by the RPE back to the circulation [57]. These lipoproteins called BrM-LLP are distinct from circulating plasma lipoproteins [92]; esterified cholesterol is the largest component of BrM-LLP, it is over 10-fold more abundant than triglyceride [93]. Our transcriptomic analyses highlights modulation of numerous genes associated with lipoprotein metabolism including, *ApoO*, *Pltp*, *Mttp*, *ApoC4* and *ApoA2* all of which are downregulated. As a group these genes are associated with plasma lipoprotein assembly, remodeling, and clearance. Of note, *Mttp*, encoding for microsomal transfer protein is necessary for the synthesis of β-lipoproteins, likely BrM-LLP by the RPE [56,92,94]. Interestingly, two genes, *ApoBec1* and *ApoBec3*,involved in apolipoprotein B mRNA editing are both upregulated. What role this gene editing activity plays in RPE-specific β-lipoproteins assembly is unknown. Further studies are necessary to understand how cholesterol biosynthesis and packaging in lipoproteins is influenced by LAP with an emphasis on therapeutics designed to restore lipid homeostasis [58].

The metabolic fate of tissues is not only regulated by the levels of rate-limiting metabolic enzymes but also by the expression and localization of solute transporters. Glucose is the primary metabolic substrate of the neural retina and is transported from the choroidal blood supply to the outer retina by GLUT1 (*Slc2al*) transporters in the apical and basolateral membranes of the RPE. The lactate generated in the outer retina through aerobic glycolysis is transported out of the retina by lactate transporters expressed in the apical (MCT1) and basolateral (MCT3) membranes of the RPE. We found the genes encoding these transporters (*Slc2a1*, *Slc16a1* and Slc16a8) were all decreased in the *LC3b*^-/-^ RPE (Figure 6A). Previous studies have shown MCT3 expression is reduced by mitochondrial dysfunction and RPE atrophy [95]. Furthermore, *SLC16a8* (MCT3) is a risk allele for AMD and loss of MCT3 expression was observed at early stages of AMD [96,97]. The reduced MCT3 expression in the *LC3b*^-/-^ RPE is consistent with previous findings showing that expression of RPE signature genes are impacted by the mitochondrial dysregulation and inflammation [98].

### 3.2. Defective LAP, inflammation and complement system

Progress in understanding of LAP in professional and specialized phagocytes has shown some interesting parallels. In macrophages and other immune cells, defective LAP leads to increased inflammatory signaling and inflammatory cytokine production [3,99,100]. RPE cells are one of the “most oxidative environments” in the body [101,102], generating reactive oxygen species resulting from an oxygen rich environment, high metabolic activity, daily flux of polyunsaturated fatty acids due to OS phagocytosis, and bisretinoid accumulation [6,103,104]. Chronic oxidative stress contributes to numerous retinal degenerative diseases including AMD [105,106]. In the absence of LC3b, undigested lipids act as substrates for lipid peroxidation reactions, there is accumulation of the proinflammatory sterol, 7-ketocholesterol (7KCh), reduced biosynthesis of anti-inflammatory molecules – NPD1 and maresin-1, thus further exacerbating inflammation [24,28,107,108]. Our GSEA analyses show that inflammatory and complement pathways are significantly upregulated in the *LC3b*^-/-^ potentially contributing to AMD like phenotype. Polymorphisms in complement factor genes in the context of environmental risk factors are associated with AMD [109–111]. Several studies point to intimate relationship between oxidative stress, complement activation, and inflammation in models of AMD [57,109,112–115]. Moreover, in an in vitro RPE lipid steatosis model, in which loss of the autophagy protein, LC3b, results in defective phagosome degradation and metabolic dysregulation, we show that lipid overload results in increased gasdermin cleavage, IL-1β release, lipid accumulation and decreased oxidative capacity [90]. The current RNAseq analyses expands these in vitro studies, as numerous cytokines and caspases (inflammatory response) are associated with the GSEA leading edge gene subsets in *LC3b*^-/-^ RPE (See Figure 5A and S1A)

### 3.3. Lapses in LAP and AMD associated phenotype

Lipid metabolic imbalance is a hallmark of cardiovascular diseases, type 2 diabetes, Alzheimer’s disease and AMD [116–122]. Moreover, several age-related disorders, including neurodegenerative disorders and AMD are associated with dysregulated autophagy [123]. Defects in autophagy and associated processes in long-lived post-mitotic cells like RPE that are challenged with lipid rich OS throughout life makes them especially vulnerable to age-related pathologies [11,28,70]. A recent study on a cohort of Finnish patients reported several SNPs in the autophagy related genes including rs73105013-T for *MAP1LC3A* that are associated with increased risk for wet AMD [124]. With a range of structural, metabolic, and functional defects, especially with regards to lipid dysregulation and inflammation, *LC3b*^-/-^ RPE offers an opportunity to investigate AMD like phenotype in a mouse model. Using the comparative transcriptomics approach, here we identified 8 DEGs overlapping with a list of prioritized genes associated with independent genetic signals in the largest AMD GWAS published to date [38,62]. These include: *Htra1*, *Tnfrsf10b*, *Tgfbr1*, *Trpm3*, *Tspan10*, *Slc16a8*, *Adamts9*, and *Rdh5* (mouse orthologs). Four of these genes: *Tspan10*, *Tnfrsf10b*, *Htra1*, and *Rdh5* also overlapped with a list of putative causal genes for AMD based on human eye eQTL (expression quantitative trait loci) and GWAS signal co-localization [125]. There is ongoing research in the field to investigate AMD pathogenesis using a range of approaches such as, gene expression studies on post-mortem eyes from AMD patients, in vitro studies with iPSC-RPE cells from subjects carrying risk allele variants, and animal models [126]. iPSCS-RPE cells derived from a patient carrying two copies of *SLC16A8* risk allele (rs77968014-G) showed deficit in transepithelial lactate transport [127]. Furthermore, there was a progressive loss of MCT3 with increasing severity of dry AMD [96]. Increase in expression of HTRA1 protein was reported in iPSC-RPE cell line from subjects carrying high risk genotype at 10q26 locus [66,128,129] and a clinical trial evaluating inhibition of HTRA1 is ongoing [130]. Mutations in *RDH5* have been reported in a subset of patients with macular atrophy [64,131]. A variant in *RDH5* has been shown to be associated with increased skipping of exon 3, nonsense-mediated decay of the of aberrant transcript, and lower minor allele specific expression, thus providing a potential mechanistic link by which *RDH5* allele contributes to AMD risk [65]. Interestingly, expression of both *RDH5* and *TRPM3* decreases under inflammatory conditions *in-vitro*, and has been suggested to contribute to RPE dysfunction in AMD [132,133]. *TRPM3*, codes for a calcium permeable cation channel activated by noxious heat, neurosteroid pregnenolone sulfate, or osmotic pressure [132–135]. *TRPM3* gene hosts *miR-204*, a microRNA highly expressed in the RPE that plays an important role in gene regulation and retinal/RPE homeostasis [135–137]. Furthermore, cholesterol enrichment inhibits TRPM3 activation and has been suggested as a mechanism for proinflammatory cytokine secretion associated with atherosclerotic processes [138]. Modulation of *Trpm3* levels along with cholesterol accumulation and increased inflammation observed in the *LC3b*^-/-^ mouse RPE offers potential mechanistic insights into AMD pathogenesis [28]. Downregulation of *Trpm3* may also reduce inflammatory thermal hyperalgesia as has been seen in *Trpm3^-/-^* mice [139,140]. The RNAseq data using the *LC3b*^-/-^ mouse model reported here is a valuable addition to bridge the gaps in our understanding of AMD pathophysiology and to identify potential therapeutic targets; and has the potential to be extended to other lipid metabolic defects/ age-related diseases as well.

## 4. Materials and Methods

### 4.1. Animals

*LC3b*^-/-^ mice (strain name: B6;129P2-*Map1Lc3btm1Mrab*/J; stock # 009336 [69], and C57BL6/J wildtype (WT) mice were purchased from Jackson Laboratory (Bar Harbor, ME). The mouse lines were confirmed to be free of the *rd8* mutation by Transnetyx. Maintenance of mouse colonies and all experiments involving animals were as described previously [12]. Mice were housed under standard cyclic light conditions: 12-h light/12-h dark and fed ad libitum, with both female and male mice used in these studies. All procedures involving animals were approved by the Institutional Animal Care and Use Committees (IACUC) of the University of Pennsylvania and were in accordance with the Association for Research in Vision and Ophthalmology (ARVO) guidelines for use of animals in research.

### 4.2. Antibodies

Primary antibodies used were: mouse anti-β-Actin (A2228; Sigma-Aldrich, St. Louis, MO), goat anti-p-cadherin (AF761, R&D), rabbit anti-MCT3 [141], rabbit-anti-GAPDH (#5174, Cell Signaling), mouse anti-RPE65 (NB100-355, Novus Biologicals). Secondary antibodies used were: goat anti-mouse and goat anti-rabbit horseradish peroxidase (HRP)-conjugated antibodies (Thermo Fisher Scientific), donkey anti-goat and anti-rabbit IgG Alexa Fluor 594/ 488 conjugates (Invitrogen).

### 4.3. RPE cell isolation

RPE cells from WT (N=4) and the *LC3b*^-/-^ (N=3) mice (age ~24 months) eyes were isolated by enzymatic treatment and gentle dissociation as described [142] with minor modifications. Briefly, the mice were anesthetized by CO2 asphyxiation, the eyes were enucleated, muscles and connective tissue were removed in ice cold HBSS-H–(HBSS without calcium or magnesium + 10 mM HEPES). Cornea, ciliary body and lens were removed, and the eyecups were incubated in 1mg/ml Hyaluronidase solution (in HBSS-H-) at 37 °C, 5% CO2 for 20 min followed by incubation in ice-cold HBSS-H+ (HBSS with calcium and magnesium + 10 mM HEPES) and gently pulling away the neural retina. Next, the RPE/choroid was incubated in 0.25% trypsin at 37°C, 5% CO2 for 1 hr, transferred to 20% fetal bovine serum (FBS) /HBSS-H+ solution and gently shaken to detach RPE sheets, followed by centrifugation at 240 x g for 5 min to collect the RPE sheets/cells.

### 4.4. RNA isolation and library preparation

Total RNA was isolated by RNAEasy Plus Mini kit (QIAgen) per manufacturer’s instructions. RNA concentration and sample quality was determined using an Agilent bioanalyzer. Total RNA samples with a RIN value of 7.1 (+/- 0.29) and a concentration between 1.1 and 2.7 ng/μl were used. 15 ng of total RNA was used for library preparation using NEB Ultra II stranded mRNA library kit as per manufacturer’s instructions and assessed for quality. Libraries were subjected to 100 bp single read sequencing on a NovaSeq SP flow cell using a NovaSeq 6000 sequencing system by the Next-Generation Sequencing Core of the University of Pennsylvania. The raw RNAseq data has been deposited in the Gene Expression Omnibus database (GEO, NCBI) under the accession number GSE225344.

### 4.5. RNAseq data analysis

Fastq files containing raw data were imported into Salmon [143] to count hits against the transcriptome defined in Gencode vM26 (annotation built on genome assembly GRCm39) [144]. Further analysis was done using several bioconductor packages in R. The transcriptome count data was annotated and summarized to the gene level with tximeta [145], and further annotated with biomaRt [146]. Principal Component Analysis (PCA) coordinates were calculated with the bioconductor package PCAtools. The normalizations and statistical analyses were done with DESeq2 [145].

### 4.6. Enrichment Analysis

DESeq2 was used to determine the significance of differential expression between the two groups. Genes showing a fold change of ≤ −1.5 or ≥ 1.5 (Log2-transformed ratio of ≤ −0.585 or ≥ 0.585), and an adjusted p-value (p-value corrected for false discovery rate using Benjamini-Hochberg method) ≤ 0.05 were considered significantly differentially expressed genes (DEGs). The up- and downregulated DEGs were tested for enrichment of Gene Ontology (GO) pathways using Metascape [147]. Additionally, an unfiltered list of genes ranked by their DESeq2 statistic was analyzed with GSEA (Gene Set Enrichment Analysis, v4.3.2) where pre-ranked analyses against gene collections in the Molecular Signatures Database (MSigDB) were carried out [148,149]. Enrichment scores and statistics for genesets in the hallmark collection were calculated.

### 4.7. Gene expression visualization

Variance stabilized expression values (calculated with DESeq2) for DEGs were used to generate heatmaps using the ClustVis web tool [150]. Rows were clustered using correlation distance and average linkage. Volcano plot was constructed using VolcaNoseR webapp using log2Fold change on the X axis and −log10(adjusted P) on the Y axis [151,152].

### 4.8. Protein-protein interaction network analysis

STRING database (version 11.5) was used to predict protein-protein interactions network (PPI) with cutoff criterion score >0.4; interaction sources included text-mining, experiments, and curated databases [153,154].

### 4.9. Immunoblotting

Immunoblotting was performed as described previously [28]. Cleared RPE and retinal lysates were prepared in RIPA buffer with 1% protease inhibitor mixture (Sigma; P8340) and 2% phosphatase inhibitor mixture 2 (Sigma; P5726). 10-15 μg of protein was separated on 12% Bis-Tris-PAGE (Invitrogen) under reducing conditions and transferred to PVDF membranes (Millipore, Billerica, MA). Membranes were blocked with 5% milk in PBS, 0.1% Tween-20 for 1h at room temperature and incubated with primary antibodies for anti-MCT3 (1:5,000), anti-GAPDH (1:10,000), anti-RPE65 (1:1000), or anti-β-actin (1:5,000) overnight at 4°C. Membranes were washed and incubated with goat anti-rabbit (1:3,000) or goat anti-mouse (1:3,000) HRP-conjugated secondary antibodies for 1h at room temperature. The blots were developed using ECL SuperSignal^®^ West Dura extended duration substrate (Thermo Scientific) and captured on Odyssey Fc (Licor) and quantified as described [35].

### 4.10. Immunostaining

Immunostaining was performed on RPE/choroid flat mounts [28]. ~24-month-old WT and *LC3b*^-/-^ mice were anesthetized, the eyes were enucleated, and incised just below the ora serrata. RPE/choroid flat-mounts were prepared by separating the retina from the RPE followed by fixation in 4% PFA for 30 min at room temp. Flat-mounts were permeabilized and blocked in blocking solution containing 5% BSA in PBS + 0.2% Triton X-100 (PBST) at 37 °C for 1 h, incubated with primary antibody diluted in blocking solution (1:2000 for rabbit anti-MCT3 or 1:200 for goat P-cadherin) at 4°C overnight, washed three times with PBST, incubated in appropriate secondary antibodies conjugated to Alexa Fluor dyes (Invitrogen, Donkey anti-rabbit 488 + donkey anti-goat 594; 1:1,000) and Hoechst 33258 (1:10,000) at 37 °C for 1 h and washed three times. Flat-mounts were mounted in Prolong Gold (Invitrogen). Images were captured on a Nikon A1R laser scanning confocal microscope with a PLAN APO VC 60× water (NA 1.2) objective at 18 °C, and the data were analyzed using Nikon Elements AR 4.30.01 software.

### 4.11. Statistical analysis

Gene level statistical tests were done using DESeq2 and pathway enrichment statistical tests were done using GSEA and Metascape. List overlaps were analyzed using the Chi squared (χ^2^) test of independence using GraphPad. A 2-tailed p-value was computed.

## Supplementary Materials

Table S1: List of differentially expressed genes (DEGs); Figure S1: String profiles for the upregulated DEGs responsible for the core enrichment of the hallmark pathways.; Figure S2: String profiles for the downregulated DEGs responsible for the core enrichment of the hallmark pathways; Figure S3: RPE signature genes.

## Author Contributions

Conceptualization, K.B-B; methodology, A.D. and J.W.T.; formal analysis, A.D., J.W.T. and K.B-B.; investigation, K.B-B. and N.J.P; data curation, A.D. and J.W.T.; writing–original draft preparation, A.D.; writing–review and editing, J.W.T., K.B-B. and N.J.P; supervision, K.B-B.; project administration, K.B-B. and N.J.P.; funding acquisition, K.B-B and N.J.P.. All authors have read and agreed to the published version of the manuscript.

## Institutional Review Board Statement

All procedures involving animals were approved by the Institutional Animal Care and Use Committees (IACUC) of the University of Pennsylvania and were in accordance with the Association for Research in Vision and Ophthalmology (ARVO) guidelines for use of animals in research.

## Data Availability Statement

The raw RNAseq data has been deposited in the Gene Expression Omnibus database (GEO, NCBI) under the accession number GSE225344.

## Acknowledgments

We thank Penn Genomics Analysis Core for support with RNAseq analyses, and PDM-Live Cell Imaging Core (University of Pennsylvania) for imaging and analyses. We thank Rachel C. Sharp and Dr Lauren Daniele for their support with the immunoblotting experiments.

