## Supplemental Figures for "Transcriptomic changes predict metabolic alterations in LC3 associated phagocytosis in aged mice"

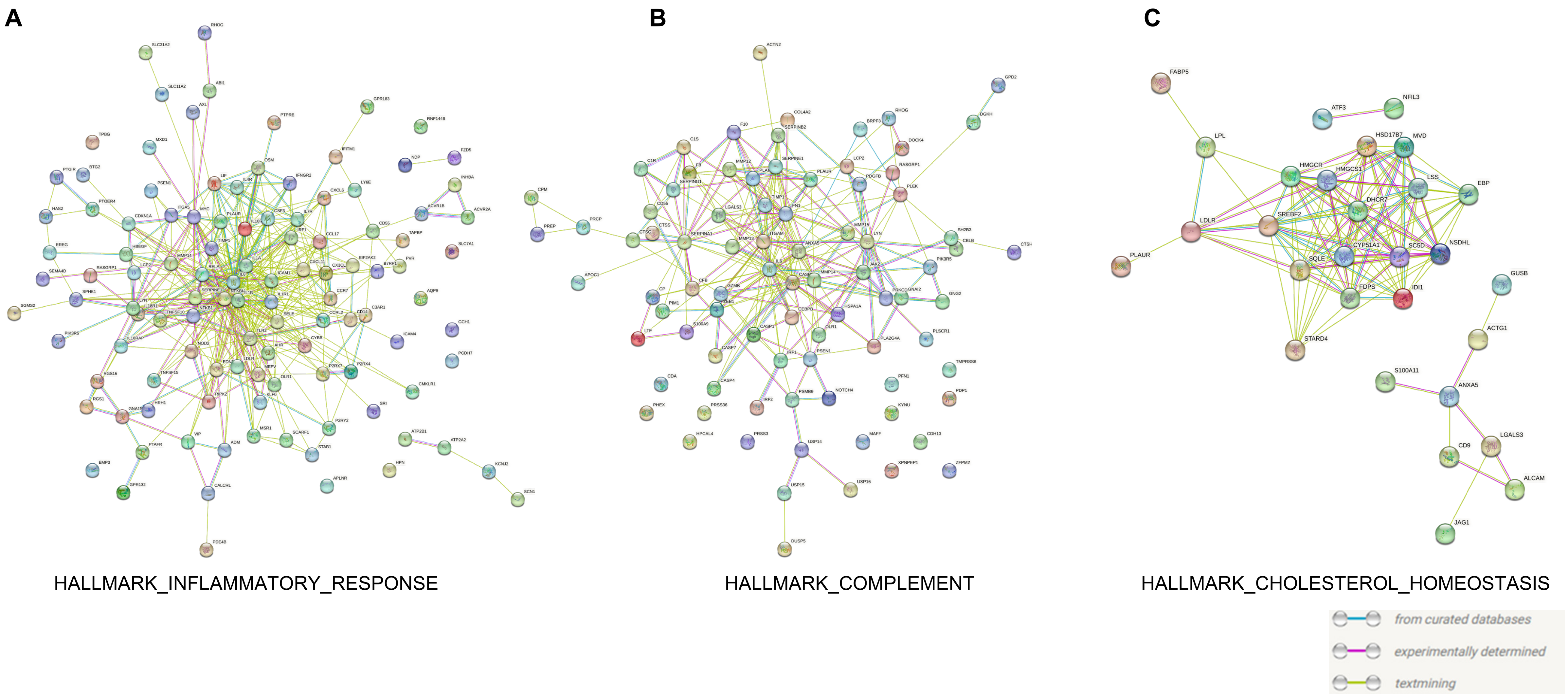


Figure S1. String profiles for the upregulated DEGs responsible for the core enrichment of the hallmark pathways.

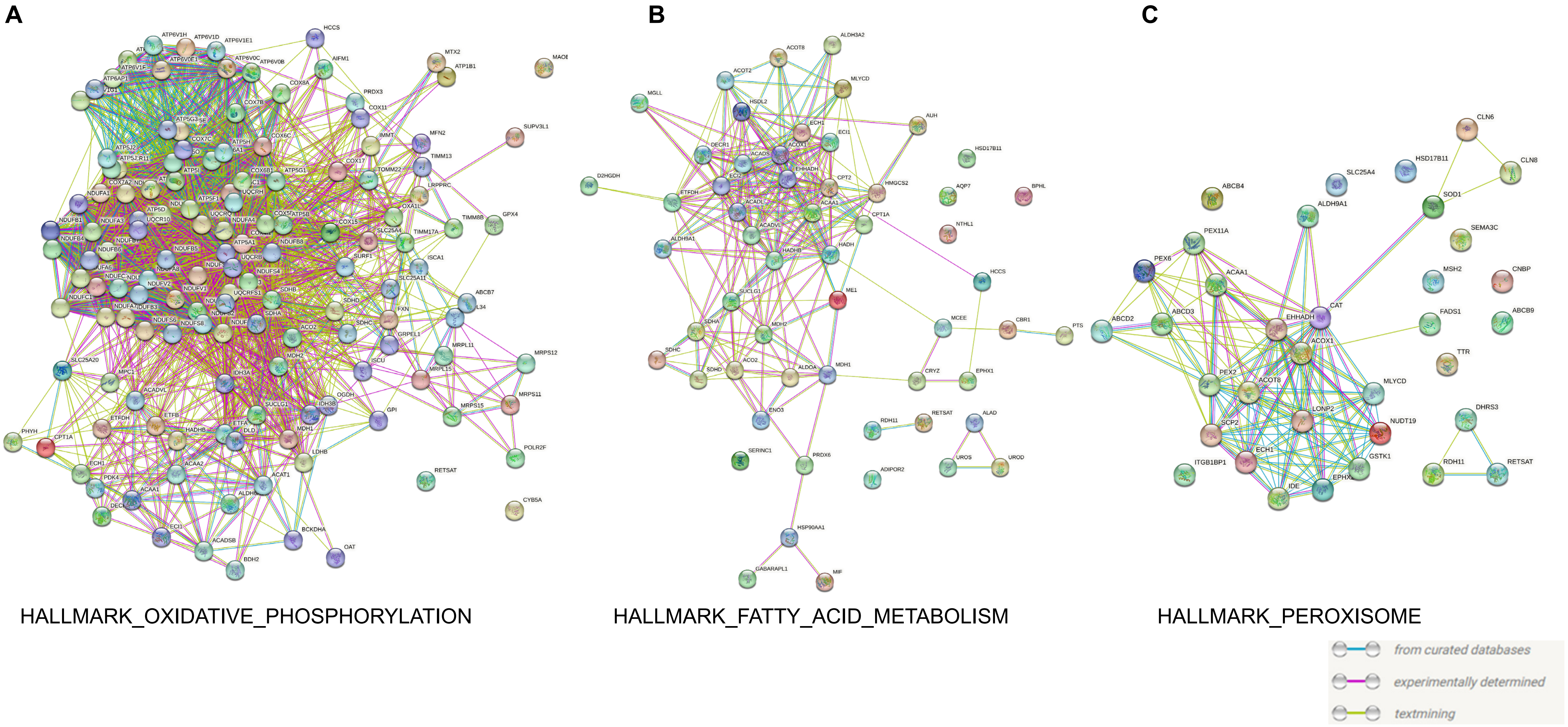


Figure S2. String profiles for the downregulated DEGs responsible for the core enrichment of the hallmark pathways.

A

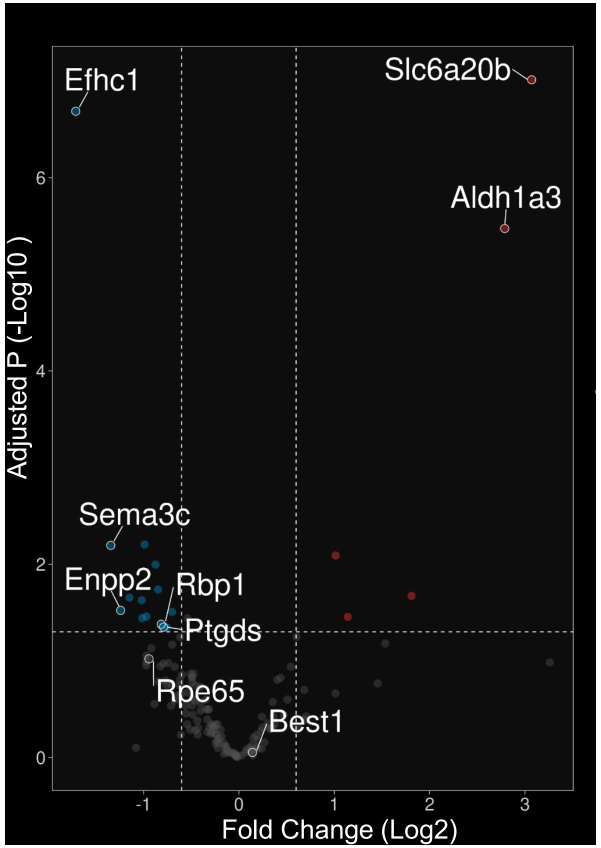


B

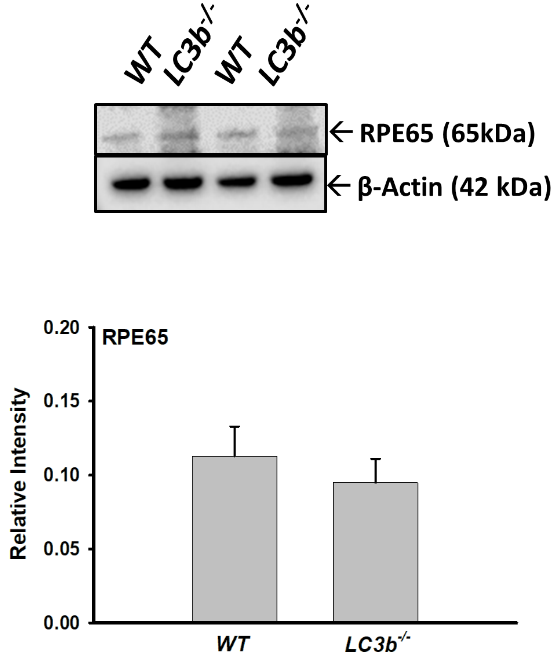


Figure S3. RPE signature genes. (A) Volcano plot showing comparison of expression of RPE signature genes between the *LC3b*$^{-/-}$ and WT RPE plotted as P adjusted (-Log 10) on the Y axis and fold change (Log2) on the X axis. Each point represents a gene. Genes that are upregulated in the *Lc3b*$^{-/-}$ are on the right and downregulated genes are on the left. DEGs were identified by using a cut-off of Padj < 0.05 and fold change of ≤ -1.5 or ≥ 1.5 and have been shown in red (upregulated DEGs) or cyan (dowregulated DEGs); grey points (unchanged genes). Top 3 genes and a selected few have been indicated. (B) *(Top):* Western blots of RPE/choroid from WT and *LC3b*$^{-/-}$. Immunoblot analysis was performed with antibodies against RPE65 and loading control (β-actin). *(Bottom)*: Quantification of mean intensity ±SEM of RPE65 relative to β-actin (N=3).
